# Concurrent prediction of RNA secondary structures with pseudoknots and local 3D motifs in an Integer Programming framework

**DOI:** 10.1101/2023.03.09.531928

**Authors:** Gabriel Loyer, Vladimir Reinharz

## Abstract

**Motivation:** The prediction of RNA structure canonical base pairs from a single sequence, especially pseudoknotted ones, remains challenging in a thermodynamic models that approximates the energy of the local 3D motifs joining canonical stems. It has become more and more apparent in recent years that the structural motifs in the loops, composed of non-canonical interactions, are essential for the final shape of the molecule enabling its multiple functions. Our capacity to predict accurate 3D structures is also limited when it comes to the organization of the large intricate network of interactions that form inside those loops.

**Results:** We previously developed the integer programming framework RNAMoIP (RNA Motifs over Integer Programming) to reconcile RNA secondary structure and local 3D motif information available in databases. We further develop our model to now simultaneously predict the canonical base pairs (with pseudoknots) from base pair probability matrices with or without alignment. We benchmarked our new method over the all non-redundant RNAs below 150 nucleotides. We show that the joined prediction of canonical base pairs structure and local conserved motifs (i) improves the ratio of well-predicted interactions in the secondary structure, (ii) predicts well canonical and Wobble pairs at the location where motifs are inserted, (iii) is greatly improved with evolutionary information and (iv) non-canonical motifs at kink-turn locations.

**Availability:** The source code of the framework is available at https://gitlab.info.uqam.ca/cbe/RNAMoIP and an interactive web server at https://rnamoip.cbe.uqam.ca/

## 1 Introduction

The rise of RNA therapeutics [1, 2] is due to technical and computational advances in our understanding of RNA sequence-structure-function paradigm. While the prediction of all-atoms RNA structure from sequence still remains a challenge [3], many different approaches have allowed to reach interesting results in various facets of the problem.

Taking advantage of the hierarchical folding of RNA, the secondary structure composed of strong stems of canonical and Wobble base pairs form first [4], many efficient theoretical approaches have been developed to predict this secondary structure. Nonetheless, the most accurate and feasible models as in the ViennaRNA package [5] or RNAstructure [6] assume that there are no crossing interactions, no pseudoknots, since that assumption adds complexity and decreases accuracy in the thermodynamic parameters making it often impractical to use. Yet pseudoknots are abundant and important. To predict accurate secondary structure with them would be invaluable for the main 3D reconstruction tools that rely on that secondary structure [7].

The prediction of RNA with pseudoknots is in all generality in the nearest neighbour model NP-Hard [8]. The dynamic programming algorithm solving exactly the minimal free energy (MFE) structure problem and with the most general classes of pseudoknots is PKnots [9] with prohibitive complexity in time of *𝒪* (*n*^6^) and space of *𝒪* (*n*^4^). Instead of exact methods, heuristics have also been developed as HotKnots [10]. Reducing the set of achievable pseudoknot configurations (while keeping known important ones) combined with sparsification techniques Knotty [11] is able to achieve a time complexity of Θ(*n*^3^ + *Z*) (with in practice *Z < O*(*n*^4^)). While deep neural networks as SPOT-RNA [12] on the subjects have been published, rigorous benchmarks show that they still lack generability [13, 14]. By formulating the problem as an Integer Programming problem IPknot [15] has proven to be effective to predict general pseudoknotted secondary structures when maximising base pair probabilities generated by RNAfold [5], leveraging fast general solvers and without sacrificing possible outputs. Methods as BiokoP [16] developed a different approach by targeting Pareto fronts with Integer Programming, showing that the pseudoknotted structure can be predicted with greater accuracy when combining the MFE and MEA.

More recent work improved IPknot using clever heuristic to solve long sequences in reasonable time, combining a dynamic threshold with a linear-time approximation of the RNA folding partition [17].

Beyond the secondary structure RNA are composed of many different important interactions. The Leontis-Westhof (LF) classification [18] defines 12 types of possible base pairs, between any nucleotides. When describing the loops between the rigid stems using those non-canonical interactions, different methods have shown that conserved sub structures are present and important [19, 20, 21]. These databases of motifs can be leverage for not only more accurate structure prediction, but also to include geometric information beyond canonical and Wobble base pairs.

In previous work we used conserved structural motifs to select an optimal secondary structure and ease all-atoms 3D reconstruction [22, 23]. Subsequently a different group developed BiORSEO [24] that computes the Pareto front of an objective function balancing structures with pseudoknots and motifs insertions. They enforce stricter constraints on the motif insertions. As discussed later, interior loop and multiloop motifs are composed of non-sequential strands, which when inserted are connected by base pairs with BiORSEO. It is not the case in RNAMoIP. The size of the Pareto front becomes also prohibitive to compute at a faster pace.

In this paper we expand on our IP framework RNAMoIP [22] to achieve **the simultaneous prediction of secondary structure with pseudoknots and structural motifs insertion with or without alignments** incorporating ideas from IPknot [15], using a newly designed local structural modules dataset computed from [25].

In Sec. 2.3 the unified IP equations are presented. We discuss in Sec. 3.9 how a sequence alignment can be used, a feature implemented in our software. We benchmark the secondary structure prediction on all known non-redundant RNAs with a determined pseudoknotted structure below 150 nucleotides in Sec. 3.4, and how the predictions fare for the canonical and non-canonical interactions inside the motifs in Sec. 3.5 and 3.6. We then evaluate how a good sequence alignment can improve the prediction in Sec. 3.9 with a set of hand aligned structures by Rfam. We conclude on analysis in Sec. 3.10 by looking specifically at the kink-turn motif that was present in 4 previous structures and how using an alignment or not influences its prediction.

## 2 Methods

Let *ω* an RNA sequence, and Ω its secondary structure. A base pair (*i, j*) *∈* Ω must be canonical (G-C or A-U) or Wobble (G-U), and have *j* − *i >* 3. The Leontis-Westhof (LW) classification of RNA base pairs [18] defines 12 different geometries possible combining two edges between Watson-Crick (W), Hoogsteen (H), Sugar (S) and an orientations cis (c) or trans (t). The canonical and Wobble base pairs are all of type cis Watson-Crick/Watson-Crick (cWW). Generally any combination of nucleotides can form any type of base pair. Stability in the nearest neighbour model is obtained from stacked base pairs [26], forbidding lonely base pairs implies formally that if (*i, j*) *∈* Ω ⇒ (*i* − 1, *j* + 1) *∈* Ω or (*i* + 1, *j* − 1) *∈* Ω.

The secondary structure Ω can be decomposed in an ensemble of pseudoknot-free structures Ω^1^, …, Ω^*m*^. Ω^*q*^ is pseudoknot-free if there is no crossing between any base pairs, formally for all (*i, j*), (*k, l*) *∈* Ω^*q*^ ⇒ *i < k < l < j* or *k < i < j < l*.

The workflow of our framework is presented in Fig. 1:

**Figure 1:**
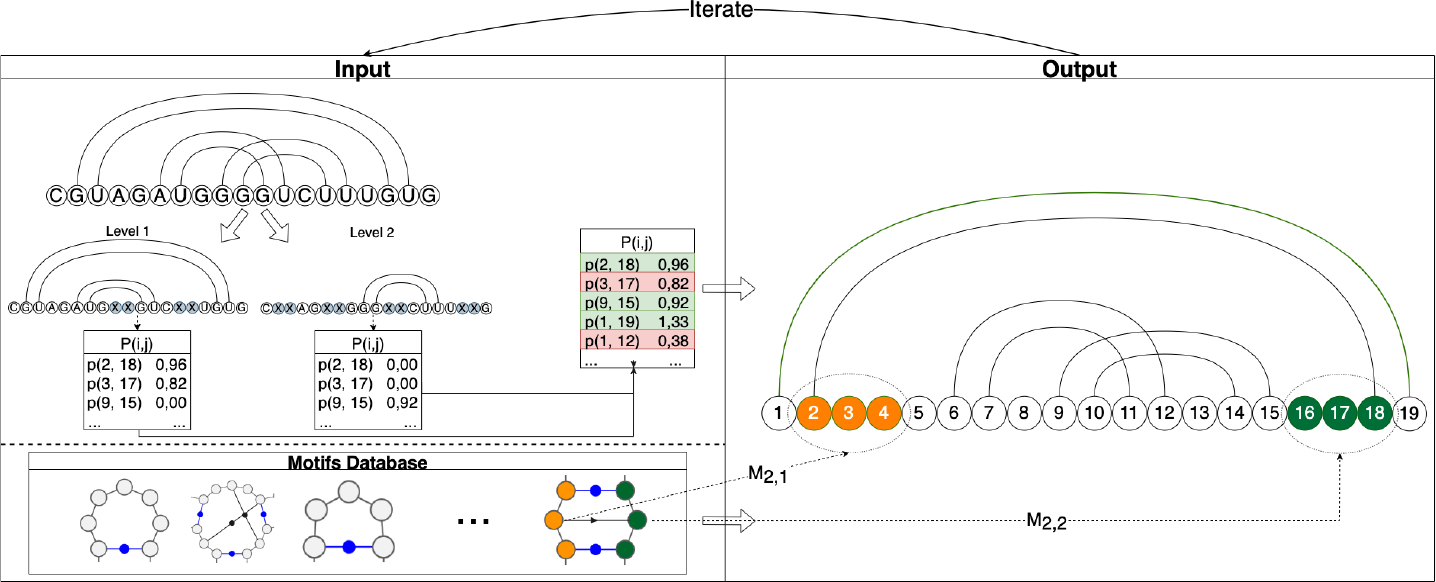
The RNAMoIP workflow. Left top: the sequence with a structure (which can be empty). The structure is decomposed in pseudoknot-free substructures, for each a constrained BPP is computed, and they are all summed together. Left bottom: a database of structural motifs containing hairpins, interior loops and bulges, and k-way junctions. Right: outputs an optimal combination between a secondary structure with pseudoknots and motifs inserted in sequence compatible locations. Each motif strand must stack or overlap by 1 position a base pair in the secondary structure.

1. Given a sequence *ω* and a secondary structure Ω simultaneously:
  a. Decomposed Ω in pseudoknot-free structures, for each compute a constrained version of a classic folding algorithm to obtain a base pairing probability matrix (if no structure is provided, the algorithm runs without constraints once), sum all the matrices.
  b. Find all possible motifs location using pattern match in the input sequence *ω*.
2. Solve the IP model to find the optimal combination of base pairs with pseudoknots and motifs given our objective function detail in Sec. 2.3.3.
3. Iterate until:
  a. Two iterations give the identical solution.
  b. A threshold in the number of iterations or time is reached.

### 2.1 Structural motifs

The validation of structure prediction using motifs database is challenging to ensure that the benchmark is not biased. We decided to use the same database as in the first version of RNAMoIP built in 2012 [22]. Additional measures to avoid overfitting are discussing in Sec. 3.3.

This database is based on the detection of similar networks of interactions among all RNA 3D structures [21]. The entire dataset is built upon 398 unique common subgraphs (or RINs - Recurrent Interaction Networks), that can be divided into 5278 different nucleotide sequences. Those sequences are composed of multiple strands, which can represent motifs like hairpins, interior loops, and multi-loops.

### 2.2 Base pair probabilities

Different tools as the ones provided by ViennaRNA [5] and RNAstructure [6] can accurately, in a thermodynamic setting, utilize the most recent set of nearest neighbor parameters to compute base pairing probabilities for pseudoknot-free structures. Following the model of IPknot [15], the secondary structure Ω is decomposed in an ensemble of pseudoknot-free structures Ω^1^, …, Ω^*m*^. For each, a base pair probability (BPP) matrix can be computed such that the base pairs in that sub-structure are enforced as hard constraints, and any position in another sub-structure are forbidden to pair. In the IP formulation, the weight of the base pair (*i, j*) will be the pseudo-probability 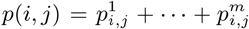 . Note that during the evaluation of the BPPs the base pairings are considered as hard constrained, they must be preserved, there is no such condition in the IP model.

### 2.3 Integer Programming Model

The integer programming model is quite complex and we reproduce here all equations for sake of completeness. Sec. 2.3.1 describes how the motifs are encoded. The Sec. 2.3.2 lists all the model variables. Then the objective function is detailed in Sec. 2.3.3. The constraints in regards to the base pairs placement are in Sec. 2.3.4 and the ones regarding the motifs insertion are in Sec. 2.3.5. **All modifications to the original equations of RNAMoIP are in green**, and the complete model is in Supp. Mat. 6.1.

#### 2.3.1 Input

We denote an RNA sequence as *ω* and Ω as a secondary structure compatible with it. *ω*_*i*_ is the nucleotide at position *i* and must be in *{A, C, G, U }*. The structure can be empty and may contain crossing interactions. We use *n* = |*ω*| as the length of the sequence.

Each motif in the database can be composed of a set of different sequence strands (e.g., an interior loop has two strands). The equation will differentiate between hairpins (1 strand), interior loops and bulges (two strands), and k-way junctions (3 or more strands). *Mot*^*j*^ is the set of motifs with *j*-strands and each is composed of its list of strands. For any motif *x ∈ Mot*, the length |*x*| represents how many nucleotides it contains. Formally, a position 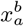 in a strand can be A, C, G, U or the wild card *, and we have:

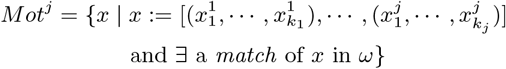

The strands are ordered in the 5^*′*^ to 3^*′*^ order of the sequence they are extracted from. The model needs to know where the *i* − *th* strand of any motif composed of *j*-parts can be inserted. These can be of different lengths, so the ensemble 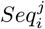 will be a set of triplets with the name of the motif and the first and last positions where the *i* − *th* strand of that motif can be inserted. The same strand can be placed in many different places. Formally:

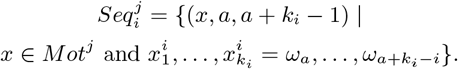

#### 2.3.2 Variables

To keep in line with the previous implementation of RNAMoIP, two type of binary variables are being tracked. First, 4 things needs to be kept track for the motifs: (1) *x* the name of the motif (2) if it is the *i*-th strand of that motif, and (3,4) the interval *k, l* where it is inserted. This will be kept done by the variable 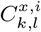 representing the insertion of the *i*-th component of the motif *x* at position (*k, l*) of the sequence. One such variable exists for every element of every set 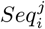. Second, for every pair of position where the BPP is above a certain threshold, the model needs to know if it is instantiated and at which level. Each level will contain a pseudoknot-free structure, and will have to be crossing a base pair in each level below it. The binary variable 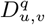 will be 1 if there is a base pair between *ω*_*u*_ and *ω*_*v*_ at level *q*. The set *ℬ* will contain all pairs of positions (*u, v*) with a potential base pair.

#### 2.3.3 Objective

Intuitively, we want to maximize the pseudo-base pair probabilities as the amount of information in the sequence. The IPknot objective function is based on the MEA and tries to maximize an approximate gain that does not take into account positions unpaired in the pseudoknotted secondary structure. With RNAMoIP we enhance the objective function by giving a bonus to unpaired positions that are known to fit into an insertable motif, compensating some of the necessary simplifications of the model. In [22], this is achieved by maximizing the square of the length of the motifs inserted, which will push to insert the smallest amount of largest motifs possible. As in Sec. 2.3.1 the number of nucleotides in motif *x* is denoted |*x*|. For each potential base pair between positions (*u, v*) the pseudo-BPPs probability *p*(*u, v*) (see Sec. 2.2) is used. As in the first version of RNAMoIP, the factor of 10 is added to normalize the scale between the pairing probabilities and the motifs insertion, which allows the *α* parameter to be calibrated more easily. A parameter *α* will be used to combine the pseudo-BPPs maximisation and motifs insertion. A weight *β*^*q*^ is used to balance the different level of pseudoknots. Following IPknot we select *β*^1^ = 0.5, *β*^2^ = 0.25, *β*^3^ = 0.125 and *β*^4^ = 0.0625. Formally, the objective function is:

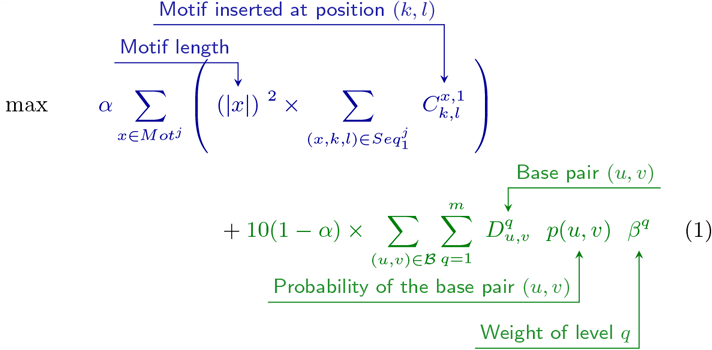

#### 2.3.4 Base pairs constraints

The first three equations will ensure that each position is only in one base pair (Eq. 2), that each level *q* contains a pseudoknot-free structure (Eq. 3), and that base pairs are added to a level if and only if they cross another base pair in every lower level (Eq. 4). Two additional properties we enforce are that base pairs must be stacked on the same level (Eqs. 5 and 6) and that at least 25% of positions must be in a base pair (Eq. 7). Note that naturally, due to the weights in the objective function, most of the base pairs will concentrate on lower levels.

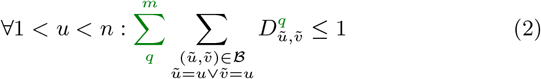

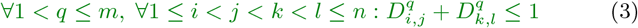

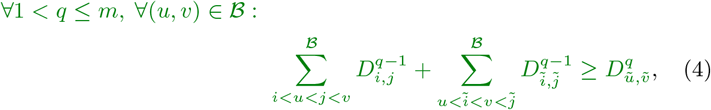

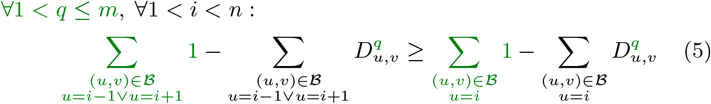

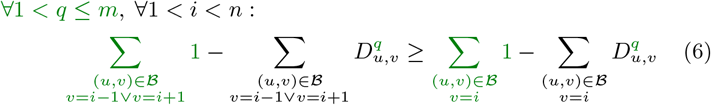

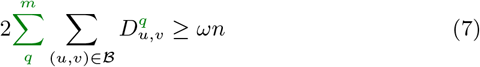

#### 2.3.5 Motifs constraints

Following the original formulation of RNAMoIP, we reproduce here all the equations necessary for the insertion of motifs. While most of them are exactly similar, there is a notable difference. One of the main conditions for the insertion of any strand is that it must be stacked or overlapping the last base pair of a strand. Since the motifs database is defined over the loops of a pseudoknot-free secondary structure, only the base pairs in the first level are considered. We give a brief overview of each equation but more details can be found in [22].

Finally, to unify both models, it is important to avoid clashes between the motifs and base pairs at the different levels. This is the role of Eq. 8. Since the structural motifs are defined on the secondary structure, we insert them in relation to the base pairs in *D*^1^ and forbid other base pairs to form inside them. We assume that motifs are a cohesive geometric unit.

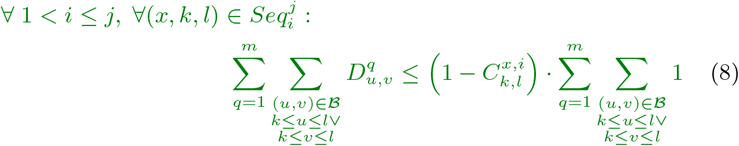

### 2.4 Incorporating evolutionary information from sequence alignments

Sequence alignment contain large amount of evolutionary information that can be leveraged for better prediction. When an alignment is provided to RNAMoIP the execution logic is slightly adapted in two ways to take this new data into account.

First, instead of relying on RNAfold, the base pairings probabilities matrix is calculated taking the alignment into account using RNAalifold [27] which is part of the ViennaRNA Package.

Second, each motif insertion score is weighted in function of its compatibility with the alignment. A motif can now be inserted if it (a) perfectly matches the input sequence at that position, or (b) is at most at Hamming distance 1 of at least 50% of the alignment at these positions. In the objective function (Eq. 1) the term 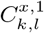 representing motif *x* inserted in position *k, l* is now weighted by the sum for each strand of the fraction of its match with the alignment. Details are shown in Sup. Mat. 6.4.

## 3 Results

### 3.1 Implementation

The Integer Programming framework is implemented in Python 3, with an interface to facilitate usage with different solvers. In this study we used the open-source solver Or-Tools [28], giving better performance. Instructions are also provided to use the open source IP solver CBC [29] through the MIP library [30], or the proprietary Gurobi [31] solver. We ran our benchmarks on Ubuntu 21.04 on an Intel Xeon Processor W-2295 with 512GB 8x64GB DDR4 2933. The source code, data, and results, are available at https://gitlab.info.uqam.ca/cbe/RNAMoIP.

### 3.2 Dataset

For benchmarking, all RNA structures between 20 and 150 nucleotides in the PDB [32] were selected, filtering for identical sequences. To avoid molecular redundancies, we kept one structure per non-redundant class as defined by the BGSU RNA Structure Atlas [33] v3.208.

The canonical base pairs in the secondary structure can be deconvoluted in different ways into a main knot-free structure and an ensemble of pseudoknots of increasing complexity [34]. A reference secondary structure from which pseudoknots are defined was determined using RNApdbee [35]. The benchmark dataset is composed of the remaining 101 structures with at least a single pseudoknot.

### 3.3 Solver

Version 2.6.4 of ViennaRNA is used to compute the base pair probability matrices. The terminating conditions are set to a maximum of 3 iterations, or two iterations with the same results. A time of 10^4^*s* was allotted for each prediction and sequences. While many optimal solutions can exist for one IP formulation only the first one achieved was used.

To avoid overfitting, before each sequence prediction all motifs belonging to any structure in the same RNA Structure Atlas v3.208 non-redundant class were removed [33].

### 3.4 Motifs insertion improve secondary structure prediction

To evaluate the capacity of RNAMoIP to predict the secondary structure, the set of True Positives (TP) consists of canonical and Wobble base pairs. The Positive Predicted Value 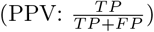, sensitivity 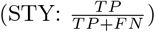 and 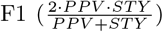are used as metrics and shown in Fig. 2. We compare the results of RNAfold and RNAMoIP with different values of *α*. When *α* = 0, no motifs are considered, and the model is equivalent to IP-knot. Note that RNAfold STY cannot reach 1, since only pseudoknotted structures are in the benchmark set, but RNAfold cannot predict crossing interactions. And this is what we observe, where it has lower STY than almost all values of *α*. On the other hand, RNAfold predictions are in general more sensitive than the IP model, especially when no motifs are used. The average values over all finished models are shown in Table 1. The optimal F1, balancing the amount of base pairs predicted and their sensitivity, is achieved with *α* = 0.15, complementing the base pairs with motifs information.

**Table 1:** Predictions results summary.

| | RNAfold | $\alpha = 0$ | $\alpha = 0.05$ | $\alpha = 0.1$ | $\alpha = 0.15$ |
| --- | --- | --- | --- | --- | --- |
| PPV | 0.63 | 0.57 | 0.621 | 0.652 | **0.671** |
| STY | 0.598 | **0.683** | 0.667 | 0.629 | 0.626 |
| F1 | 0.607 | 0.616 | 0.638 | 0.636 | **0.642** |

**Figure 2:**
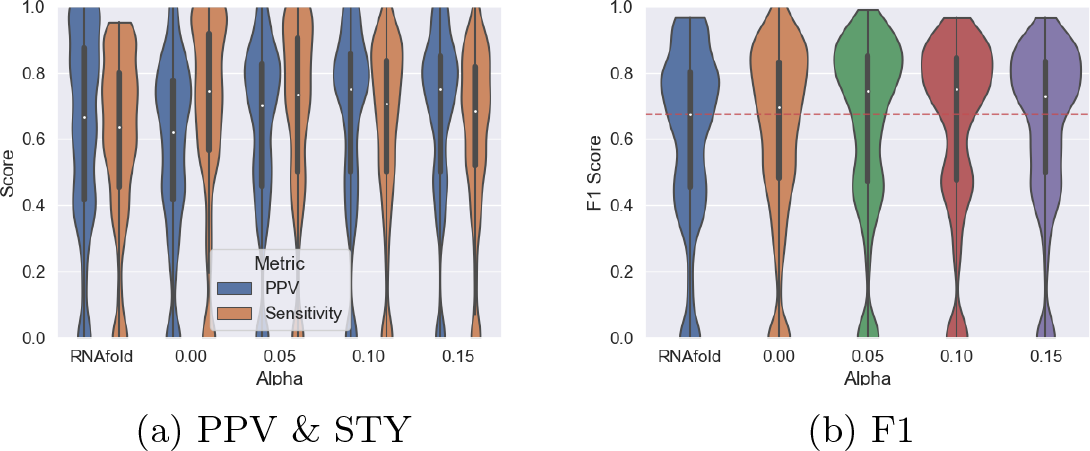
Pseudoknots prediction accuracy. Comparing results for RNAfold (cannot predict crossing interactions), without motifs insertion (*α* = 0), and for different values of *α*. When *α >* 0, all base pairs inserted in the same motifs are counted as true positives.

The statistics shown previously are computed over all the secondary structures. Our models allow up to 2 crossing levels between the base pairs yet 95 of the benchmarked structures have only 1 level of crossing interactions. No over-prediction of the pseudoknots level was observed, as shown in Table 2. Pseudoknots are in fact usually under-predicted. When no motifs are inserted, around 15% of structures have a pseudoknot level too low, up to 50% when motifs are added to the model. Nonetheless, the improvement in PPV and of over 10% in F1 measure indicates that although no pseudoknot is predicted the structure is much more accurate.

**Table 2:** Predicting pseudoknot lvl. As *α* is increased, we underestimate the lvl of pseudoknots that are present in the structure. The complexity of the IP model increases and makes it more challenging to find a feasible optimal solution in time.

| $\alpha$ | 0 | 0.05 | 0.1 | 0.15 |
| --- | --- | --- | --- | --- |
| PK lvl too low | 25 | 51 | 62 | 68 |
| PK lvl correct | 76 | 50 | 39 | 33 |

### 3.5 Recovering canonical and Wobble interactions in the motifs

For each motif inserted, we retrieved from the RNA structure atlas all canonical and Wobble interactions at the inserted positions to build our positive examples. Note that these interactions can be crossing inside the loops and do not need to be stacked, therefore they are not necessarily part of the secondary structure. Since a motif sequence in our database can match different sub-structures, the one with the best structural match was used in this and the subsequent section. The same metrics as previously, PPV, STY (available in Supp. Mat. Fig. S1), and F1 are computed and averaged over all motifs in all structures. Inside the motifs, an F1 value around 40% is achieved, a slightly lower precision for predicting the canonical and Wobble pairs in the motifs than the pseudoknotted secondary structure, as shown in Fig. 3.

**Figure 3:**
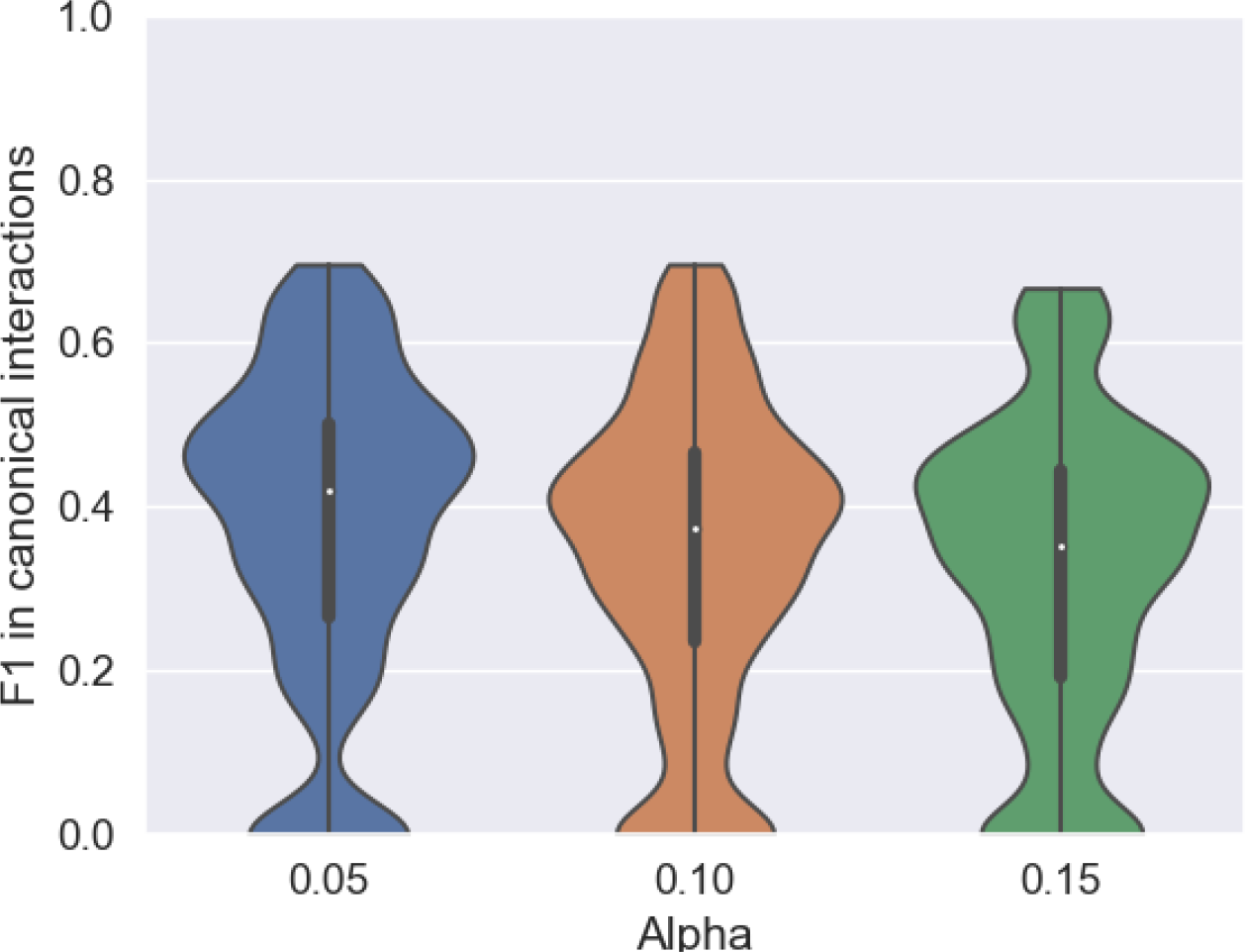
Prediction accuracy of canonical and Wobble interactions in motifs. For *α* values of 0.05, 0.1, 0.15 that more than half of the canonical and Wobble base pairs in the motifs are correctly predicted, and 40% of them are generally captured.

### 3.6 Non-canonical interactions remain challenging to predict

In the minority and hard to predict, non-canonical interactions have been shown to be necessary for many RNA functions. RNAMoIP maximizes the insertion of motifs based solely on a sequence match and the structural context. We expect similar loops in different RNA to adopt the same topology, but it is not given since each motif inserted comes from an RNA from a different redundancy class. As in the previous section, for each motif the positive set consists in the ensemble of all non-canonical interactions between inserted positions. The distribution of F1 is shown in Fig. 4 (PPV and STY in Supp. Mat. Fig. S2) highlighting how the motifs as of now are still limited to predict this finer grain information.

**Figure 4:**
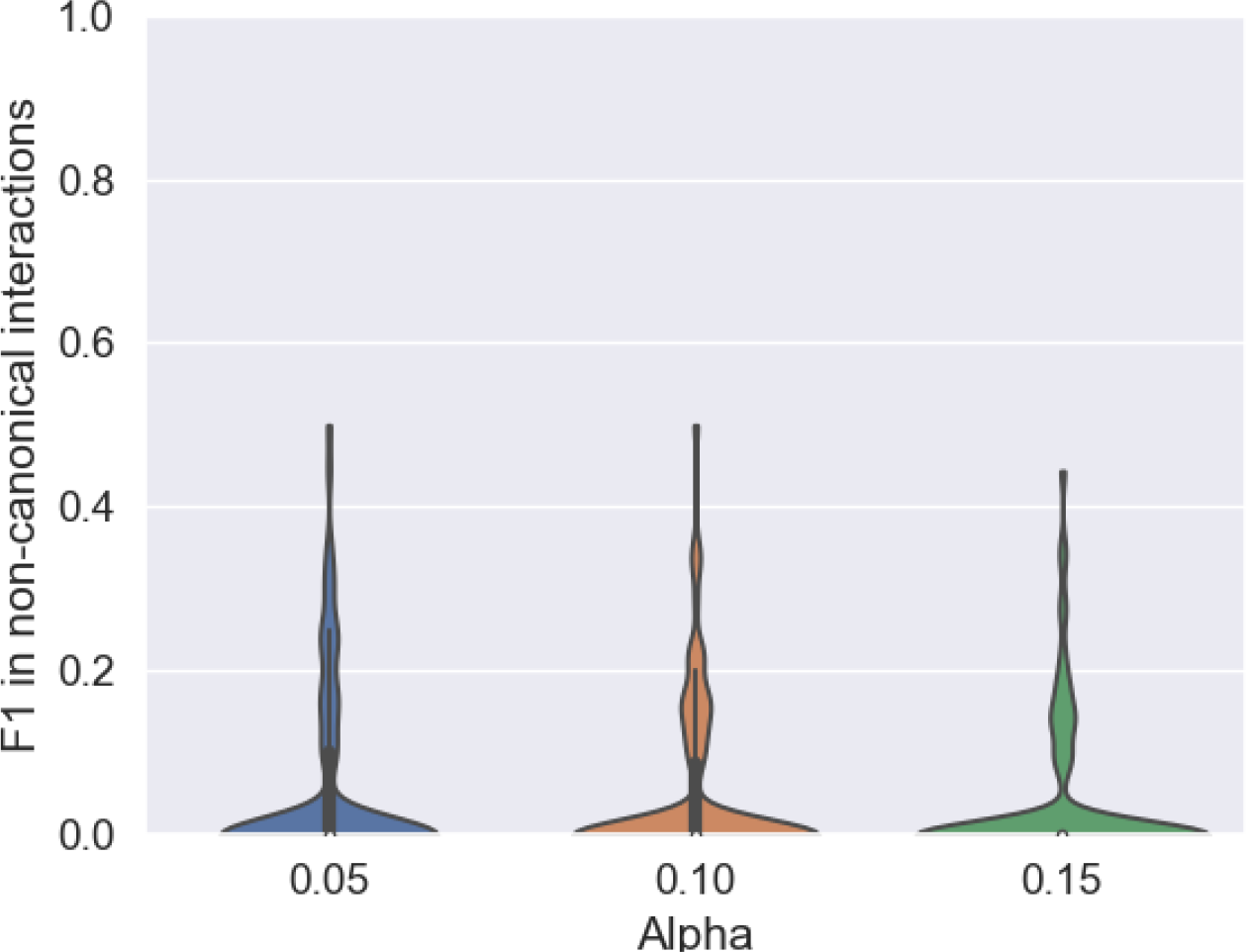
Predicting non-canonical base pairs in motifs. True positives are the non-canonical base pairs at positions where one motif is inserted in the sequence. They are composing at most 15% of the interactions in the inserted motifs, and are hard to predict (see Fig. S3 in Supp. Mat.).

While looking dramatic, it can be explained in two different ways. First, their number is really low, as shown in Fig. S3. The Y-axis shows that at these positions rarely more than 4 non-canonical interactions are present per RNA, and rarely more than 2 found in the inserted motifs. Second, this is underestimating the non-canonical interactions since many cannot be predicted by our model. On top of those that are at locations where no motifs are predicted, many can be linking motifs together, but those cannot be found in the dataset used.

### 3.7 Performance

Integer programming is known to be NP-complete, but decades of optimization have allowed to leverage efficient implementations to express complex models. We remind that a maximum of 10^4^s was enforced and the 14 sequences without a solution at the time cutoff with *α* = 0.1 were discarded. We show in Fig. S8 the execution time in seconds. More heuristics as the ones developed by IPknot for long sequences could be used, at the cost of a decrease in the accuracy. We expect that the optimal gains would be achieved by optimizing the location where we allow motifs to be inserted.

### 3.8 Tools Comparison

When benchmarked only on the canonical and Wobble base pairs, while a value of *α* = 0.15 seems to give marginally better results 1 in the secondary structures, it is a bit worse inside the motifs 3. We therefore selected *α* = 0.1 as default parameter value.

We evaluated in Fig. 5 RNAMoIP with *α* = 0.1 against 11 other tools: RNAfold [5], our implementation of IPknot [15] (i.e., *α* = 0), PKnots [9], HotKnots [10] Knotty [11], SPOT-RNA [12], MXfold2 [36], LinearFold [37], pAliKiss [38], BiokoP [16] and BiORSEO [24]. While in the task of predicting base pairs of the secondary structure RNAMoIP clearly outperforms most of the competitors. When compared with Knotty it produces as good results while providing more information in the form of motifs inserted. Since we can not re-train SPOT-RNA, it was run on the subset of the test sequences that did not share a BGSU RNA Structure Atlas redundancy class with its training set. We show in the supplementary material the results when run over the entire set, and we can see how the overfitting greatly improves its result. As mentioned in recent papers [13, 14], if more data was available, a benchmark based on Rfam families would probably decrease further the results.

**Figure 5:**
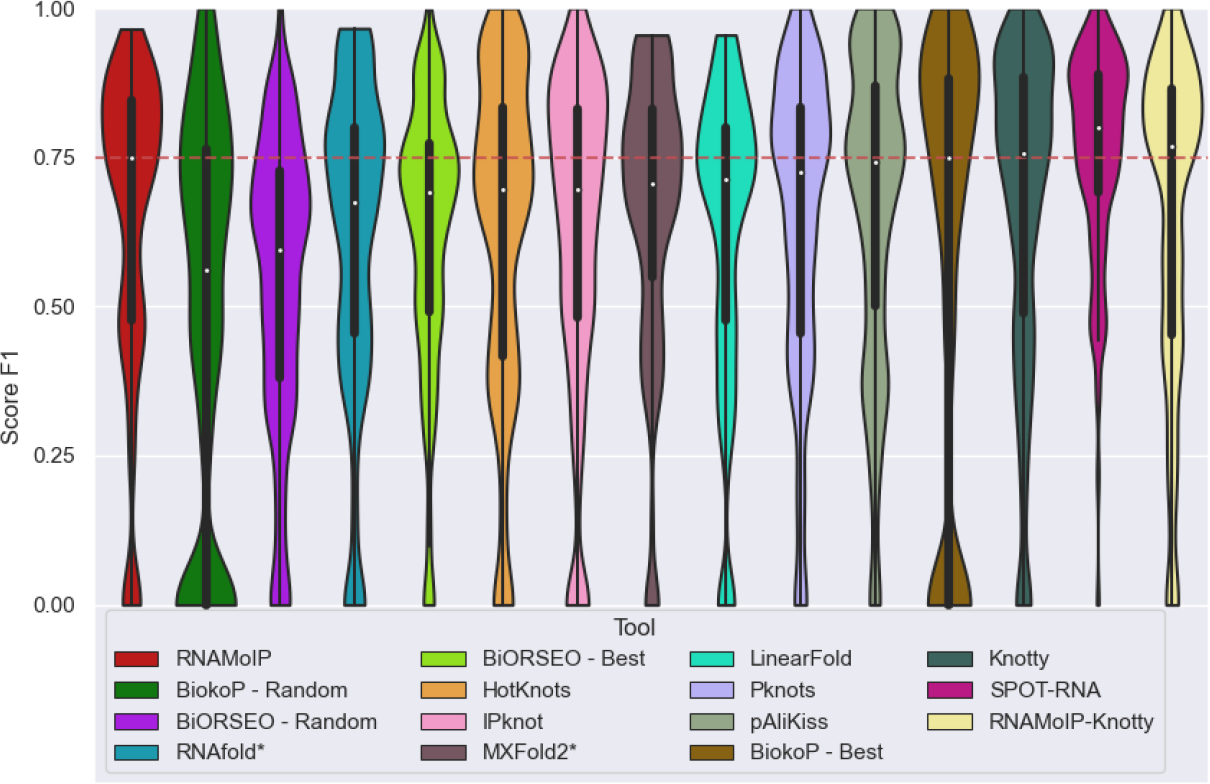
Tools comparison of F1 scores. RNAMoIP with parameter *α* = 0.1 is compared with 13 other tools. Knotty does not provide additional geometric information. SPOT-RNA is evaluated on the subset of sequences, not in its training set.

**Figure 6:**
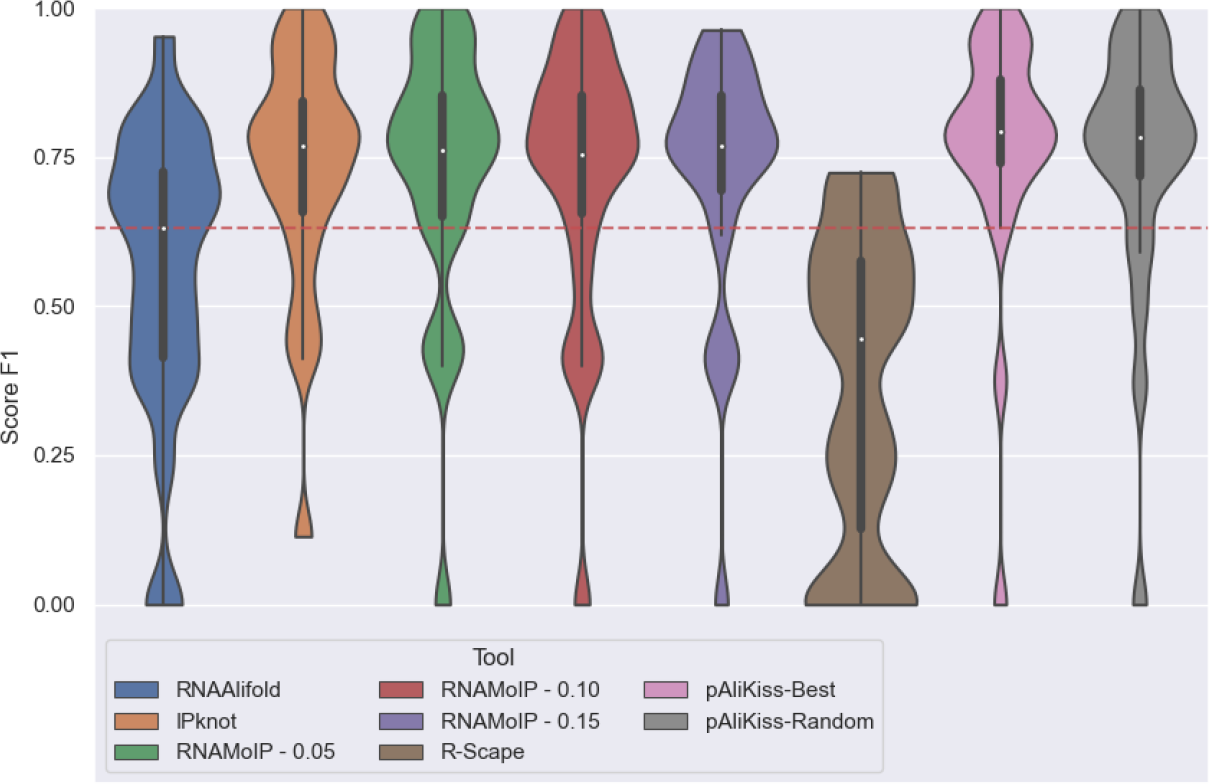
Alignment-based secondary structure prediction accuracy. Result of the alignment’s dataset with the help of the alignment informations. pAliKiss and R-scape were also added for comparison.

### 3.9 Including alignments increases predictability

Our model was extended in Sec. 2.4 to incorporate evolutionary information as sequence alignments. The Rfam database [39] is a repository of curated structured RNA families and has recently started providing hand aligned PDB structures. As of January 2023 they are 40 Rfam families alignments with 201 PDB structures aligned. Out of those, they belong to 23 different families (as defined by RNA Structure Atlas). The representative of each family was selected, or a random for the two families with a structure in an alignment but not its representative. The name of the 23 structures are provided in the directory with the code in the results folder. We describe in Supp. Mat. 6.4 how we modified the procedure.

To the best of our knowledge, only pAliKiss [38] can use an alignment to predict secondary structures with pseudoknots. Due to the limited amount of tools and data, only 23 structures, we used as baseline RNAalifold [5]—which doesn’t take into account pseudoknots—and evaluated the number of base pairs statistically supported by the alignment using R-scape [40]. We show the comparisons with RNAMoIP in Fig. S6. We compare with RNAfold and RNAMoIP on these sequences without the alignments in Fig. S5.

A drastic increase in PPV is observed as much for RNAalifold vs RNAfold as for RNAMoIP. R-scape indicates that only a fraction of the base pairs are directly supported by the sequence alignment. We see that with a sequence alignment, there isn’t a significant improvement brought by the motifs under our scheme. The F1 distributions for RNAMoIP at *α* = 0 where no motif is taken into account or *α* = 0.1 are almost similar, and increasing *α* slowly decreases the F1 value. These results are of equivalent quality to the ones returned by pAliKiss. We note that pAliKiss produced in 60% of the cases multiple solutions, up to 47, but as shown there is no observable difference between its best and a random structure. Incorporating motif prediction using our simple scheme does not decrease accuracy while providing additional information, as we will explore in the next section.

### 3.10 Kink-turn

A particularly interesting geometrically complex local motif is the kink-turn [41, 42] which is linked to many different biological processes [43]. Since they greatly constrain the structure they are key pieces completely ignored in the secondary structure representation.

The RNA Structure Atlas [20] maintains different classes of kink-turn as all their occurrences in known RNA structures. From the previous set of 23 structures with alignments, 4 have a kink-turn annotated in them all in group `IL 29549.4`, in PDB and chains 3V7E-C, 4AOB-A, 4KQY-A, and 5FJC-A. Since all four belong to the same group, they are represented with the same non-canonical diagram.

We investigate where motifs are predicted at the position of the kink turn with and without the alignment in Fig.7. In both cases, different interior loop are colored as green or blue, and hairpins as orange. When exactly the right base pair was predicted, it is also colored, the other ones inside the predicted motifs are omitted.

**Figure 7:**
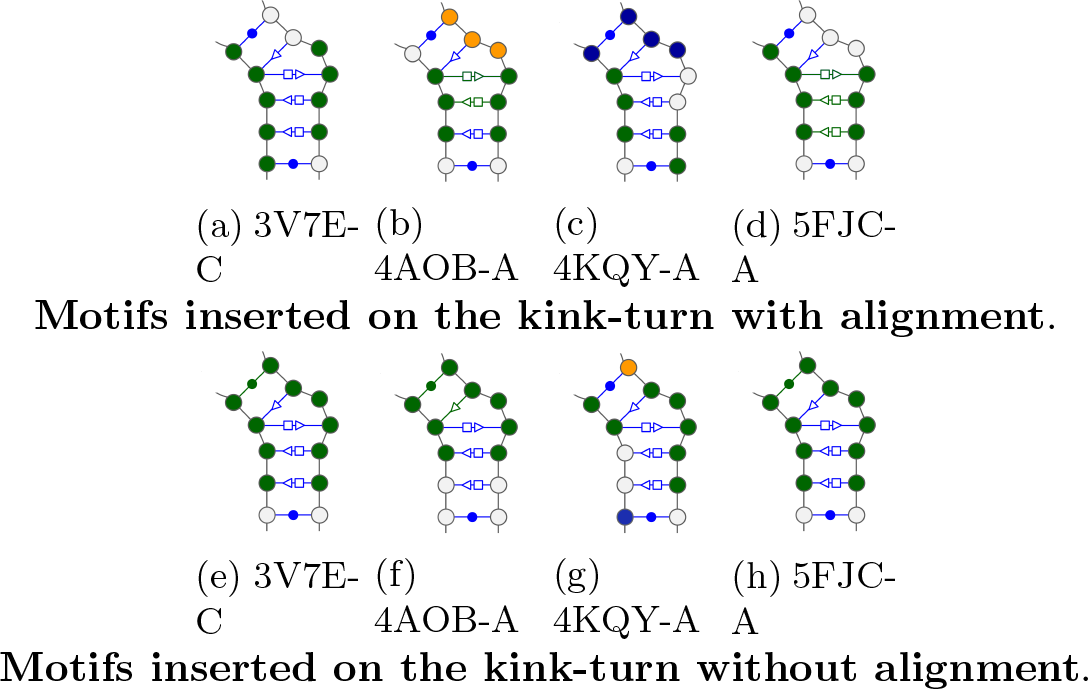
For the 4 PDB with a kink-turn the representation of it’s group and colored the position where RNAMoIP predicted motifs on the kink-turn as the exact base pair.

We observe immediately that there is always an interior loop matching on each strand of the kink-turn, in green. The prediction without alignment seem to be more cohesive, having always a match on all the position forming a cycle with the triangular pattern. But we can see using the alignment that the match on 5FJC chain A recover 3 out of the 4 non Watson–Crick/Watson–Crick base pairs. Since the non-canonical interaction do not necessitate an exact match to reflect similar geometries [44] a more in depth study out of the scope of the present work is needed. Nonetheless, RNAMoIP predicts relevant geometric information that cannot be represented by the secondary structure at the kink-turn location.

## 4 Webserver and visualisation

In an effort to make the program more accessible, a web server was developed and made available at https://rnamoip.cbe.uqam.ca. Users can submitted their own sequences and can adjust the RNAMoIP parameters, as the *α* or the maximum level of pairs crossing. After the predictions are completed asynchronously, a dashboard presents various information related to the predicted structures. The structures with the different motifs inserted are shown in a clickable 2D layout built from Varna [45]. All occurrences that correspond to each motif found are shown in their respective tab, with all their canonical and non-canonical interactions as a dynamic 3D visualisation allowing superposition of the different occurrences of the motifs.

## 5 Conclusion

In this work, an integer programming framework allowing simultaneous prediction of the secondary structure with pseudoknots, and insertion of structural motifs, is presented. The implementation in RNAMoIP is benchmarked over the 101 non-redundant pseudoknotted RNAs with known structure. We show that combining the approach of IPknot to construct pseudoknotted structures based on the base pair probability matrix obtained by the standard thermodynamic model, implemented in the ViennaRNA, with the insertion of known conserved structural motifs, allows to: (1) increase the accuracy of the prediction of secondary structure with pseudoknots, (2) generates accurate knowledge about canonical and Wobble interactions present inside structural motifs, which might not belong to the secondary structure, (3) improves drastically under a simple scheme to incorporate evolutionary information from multiple sequence alignment as validate with 23 independent structures aligned by Rfam, and (4) predict non-canonical interaction motifs at kink-turn locations.

Two main limitations are highlighted by our work. First, motifs can be inserted now based on a perfect sequence match. More advanced probabilistic techniques, as RMDetect [46], JAR3D [47] or BayesPairing [48], would allow to integrate a more rigorous term in the objective function, as match motifs with altered sequence, increasing diversity and therefore the range of predictable structure. Second, the database of motifs only incorporates loops (i.e., hairpin, interior loops, multi-loops). These approaches can directly use multiple sequence alignment which would not only increase the general base pairs as shown in this work, but would probably show a more significant boost from incorporating the motifs.

Advances in structural molecular biology are pushing against the limitation of the nearest neighbour model. While the biological importance of networks of non-canonical interactions are becoming more and more evident, the capacity to predict them lags far behind. The IP programs remains a promising direction for RNA structure determination due to the flexibility of their formulation allowing to go above the nearest neighbour model. Expending to more complex conserved structures, as groups of interacting and conserved loops containing pseudoknots described in Carnaval [21, 25] would allow to take fully advantage of the IP formulation and extend the notion of pseudoknots prediction to all non-canonical interactions. This flexible formulation will also allow to give specific rules to help incorporate chemical modifications and other features that are absent from the nearest neighbour model.

## 6 Supplementary material

### 6.1 Full Integer Programming Model

The hairpins insertion that are composed of only one strand are constrained by Eq. 9, which was modified to only consider base pairs in level 1. The insertion of interior loops and bulges must first ensure that strands are placed in acceptable positions (Eq. 10) and that the motif must fill at least 2 unpaired positions, ensuring information is added to the system (Eq. 11).

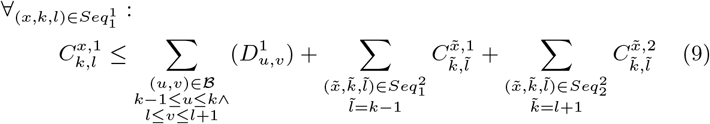

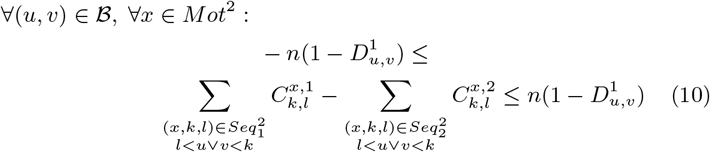

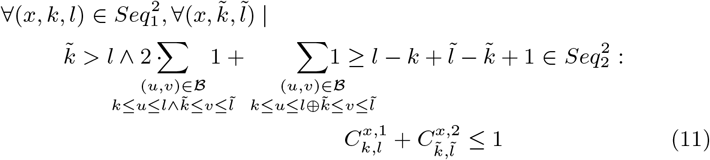

The k-way junctions admissibility of insertion is decided in Eq. 12, ensuring that each strand can be reached without crossing the base pairs in the first level. This is equivalent to Eq. 10 for the interior loops.

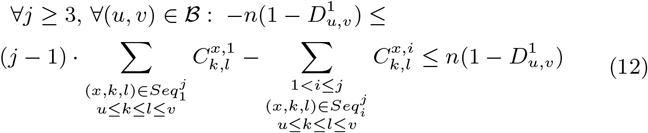

An important feature of RNA structure is that their sequence is ordered, from the 5^*′*^ to the 3^*′*^ end, and that it is not symmetric. In a motif, an order is defined over the strands following that direction. The model constrains where a strand in a motif can be placed given the insertion of the previous (Eq. 13) or next (Eq. 14) strand of the same motif. An important consideration is that at the end there must exist a mutually exclusive decomposition of the strands such that each inserted motif is complete, even if many copies are found (Eq. 15).

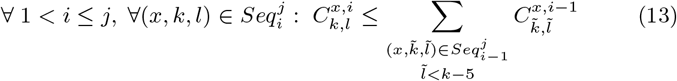

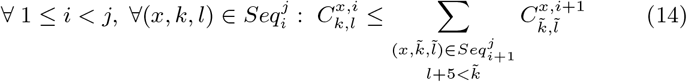

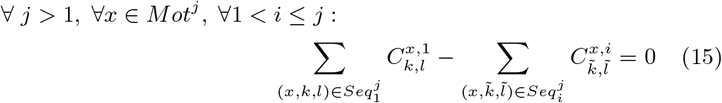

### 6.2 Predicting canonical interactions in motifs

**Figure S1:**
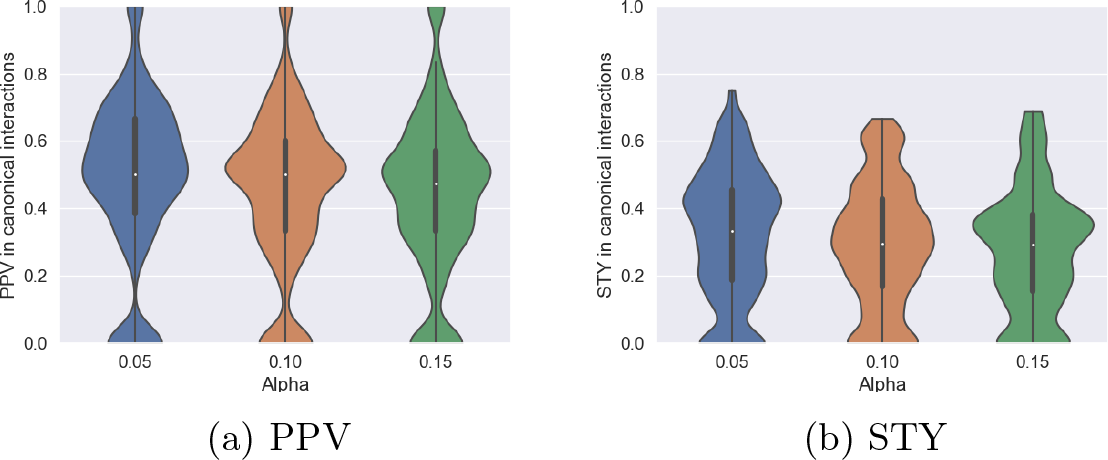
Predicting canonical and Wobble interactions in motifs. For *α* values of 0.05, 0.1, 0.15 that more than half of the canonical and Wobble base pairs in the motifs are correctly predicted, and 40% of them are generally captured.

### 6.3 Predicting non-canonical interactions in motifs

**Figure S2:**
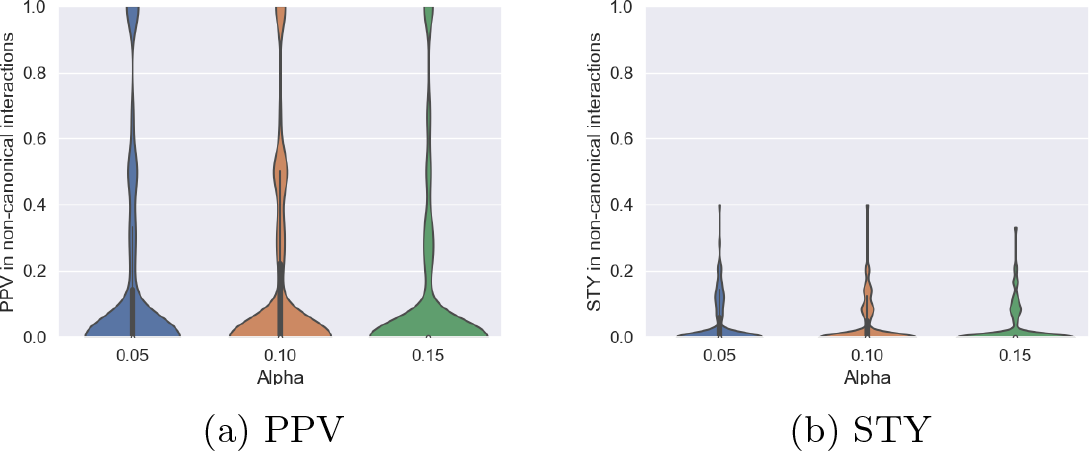
Prediction accuracy of non-canonical base pairs in motifs. True positives are the non-canonical base pairs at positions where one motif is inserted in the sequence. They are composing at most 15% of the interactions in the inserted motifs, and are hard to predict.

**Figure S3:**
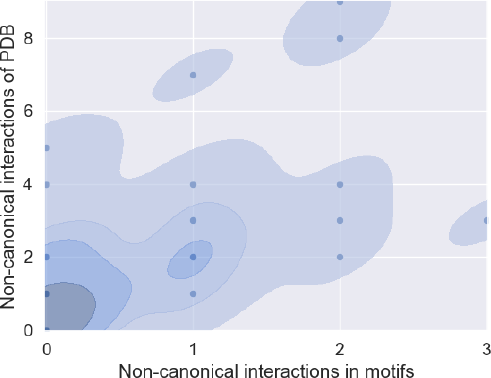
Non-canonical interactions. distribution of the number of non-canonical interactions that are observed at motifs inserted locations. On the y-axis the number in the real structure, on the x-axis how many are annotated in the inserted motif.

### 6.4 RNAMoIP on alignments

Due to the nature of the sequence alignment, we relax the procedure to insert motifs as follows. RNAMoIP predicts the structure of 1 sequence that can be enhanced with an alignment. All columns that are gaps in the sequence of interest are discarded. Therefore, for a motif component 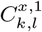 the position *k, l* are the same in the structure for which we are doing the prediction, and the alignment.

We first identify for each sequence, without gaps, positions where each motifs can fit. For each of these, we count in the alignment the fraction of other sequences that are at most at a Hamming distance of 1. If that ratio is above 50% we consider that the motif can be inserted at these positions. Formally, we define a function

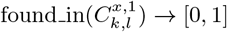

such that: found in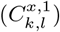 returns the fraction of subsequences in the alignment between positions *k, l* that are at most at Hamming distance one from the motif *C*^*x*,1^. Then we can have the normalizing function:

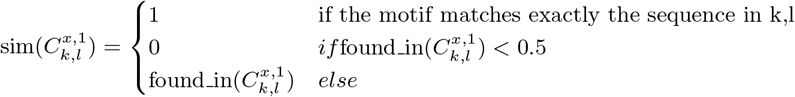

Finally the updated objective function when an alignment is provided becomes:

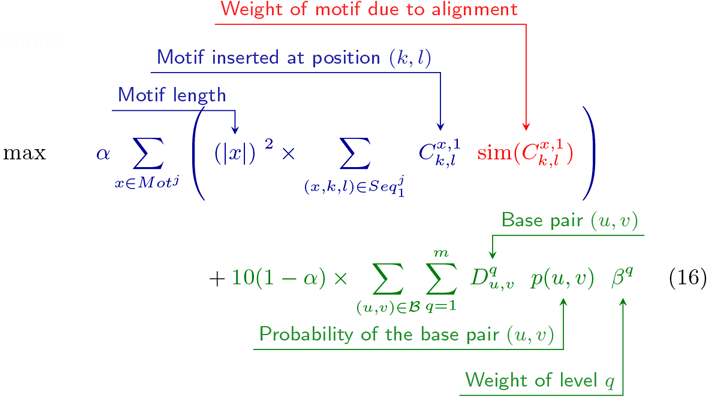

**Figure S4:**
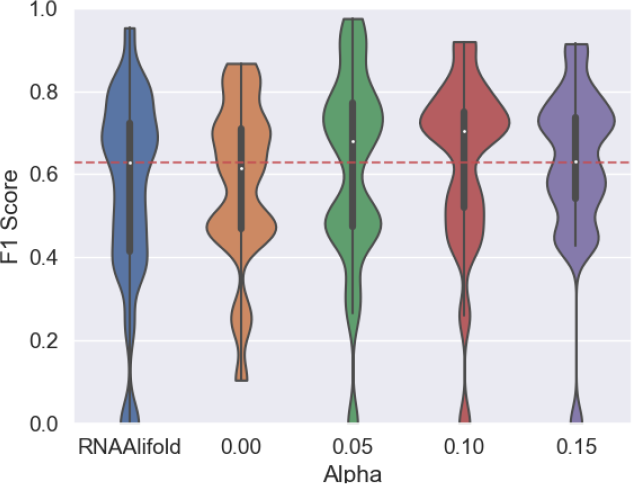
Alignment-free secondary structure prediction accuracy. Result of the alignment’s dataset without using the alignment informations.

**Figure S5:**
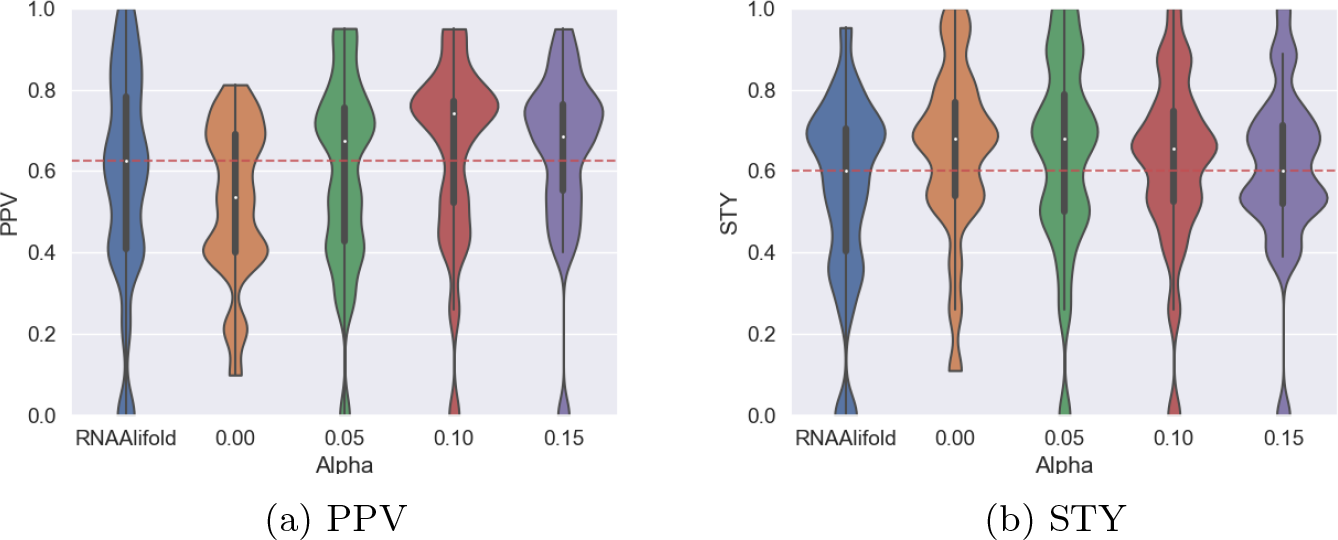
Alignment-free secondary structure prediction accuracy. Result of the alignment’s dataset without using the alignment informations.

**Figure S6:**
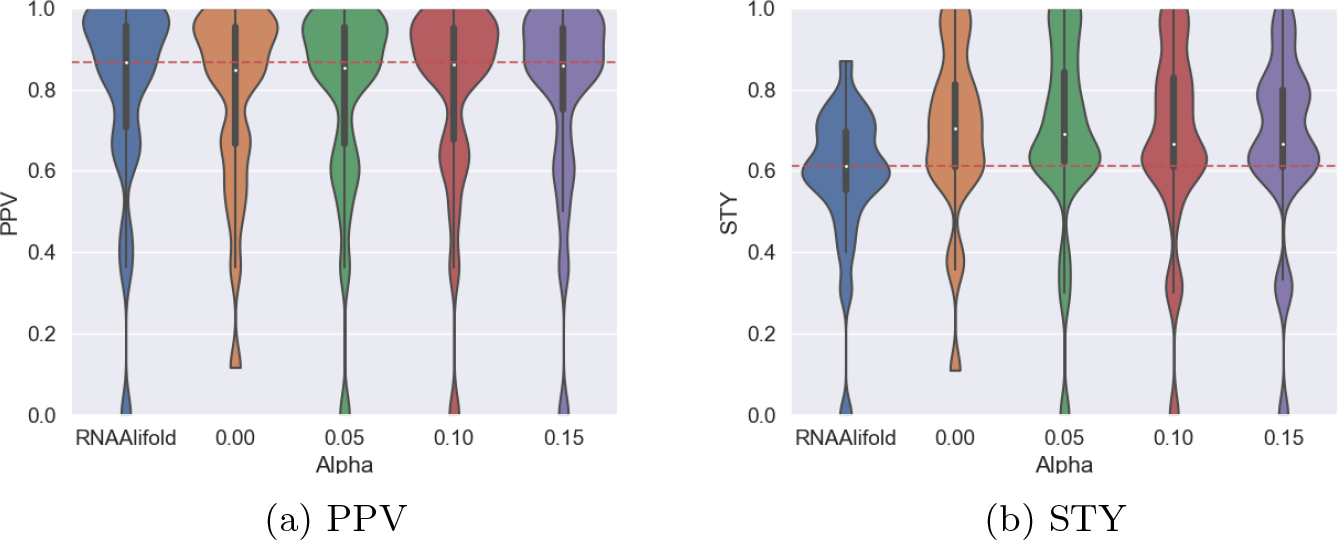
Alignment-based secondary structure prediction accuracy. Result of the alignment’s dataset with the help of the alignment informations.

### 6.5 Complete tool analysis

**Figure S7:**
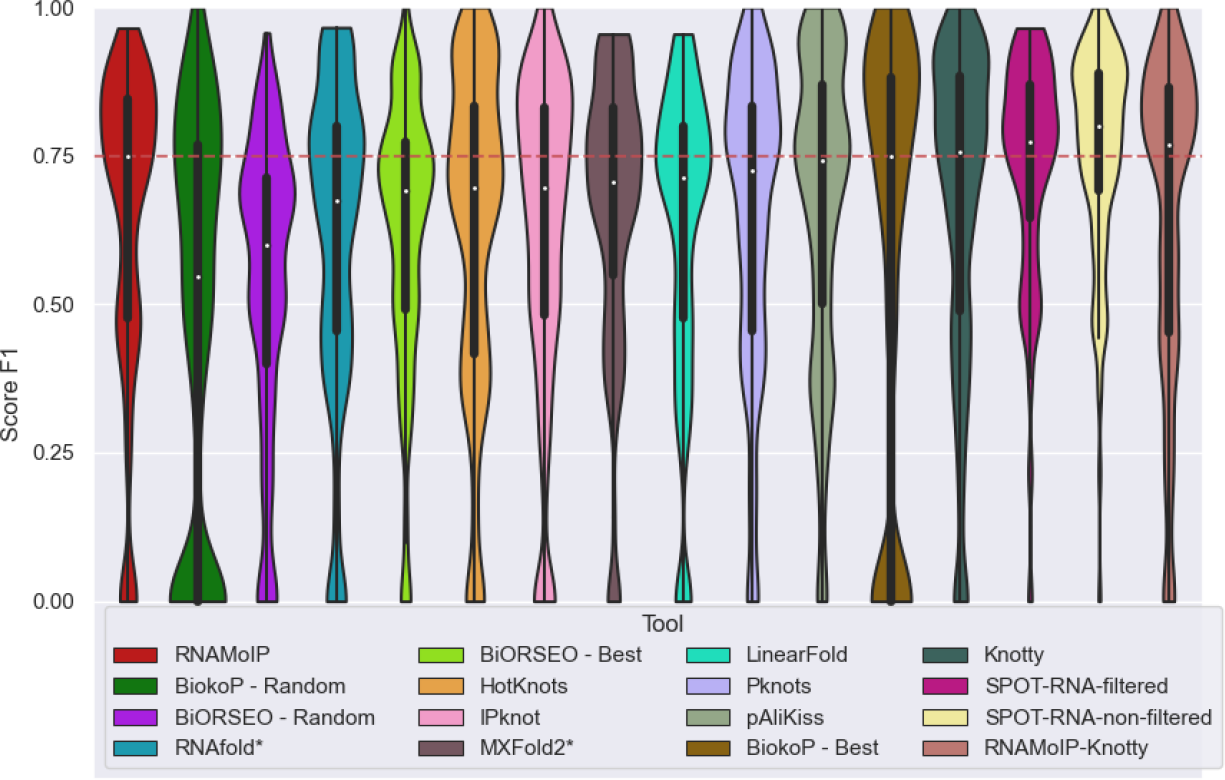
Tools comparison of F1 scores. Include two versions of SPOT-RNA. As in Fig 6 SPOT-RNA is evaluated on the subset of sequences not in its training set. We also show the results on the entire dataset, highlight the overfitting if not careful in the separation of test and train set.

### 6.6 Computation time benchmark

**Figure S8:**
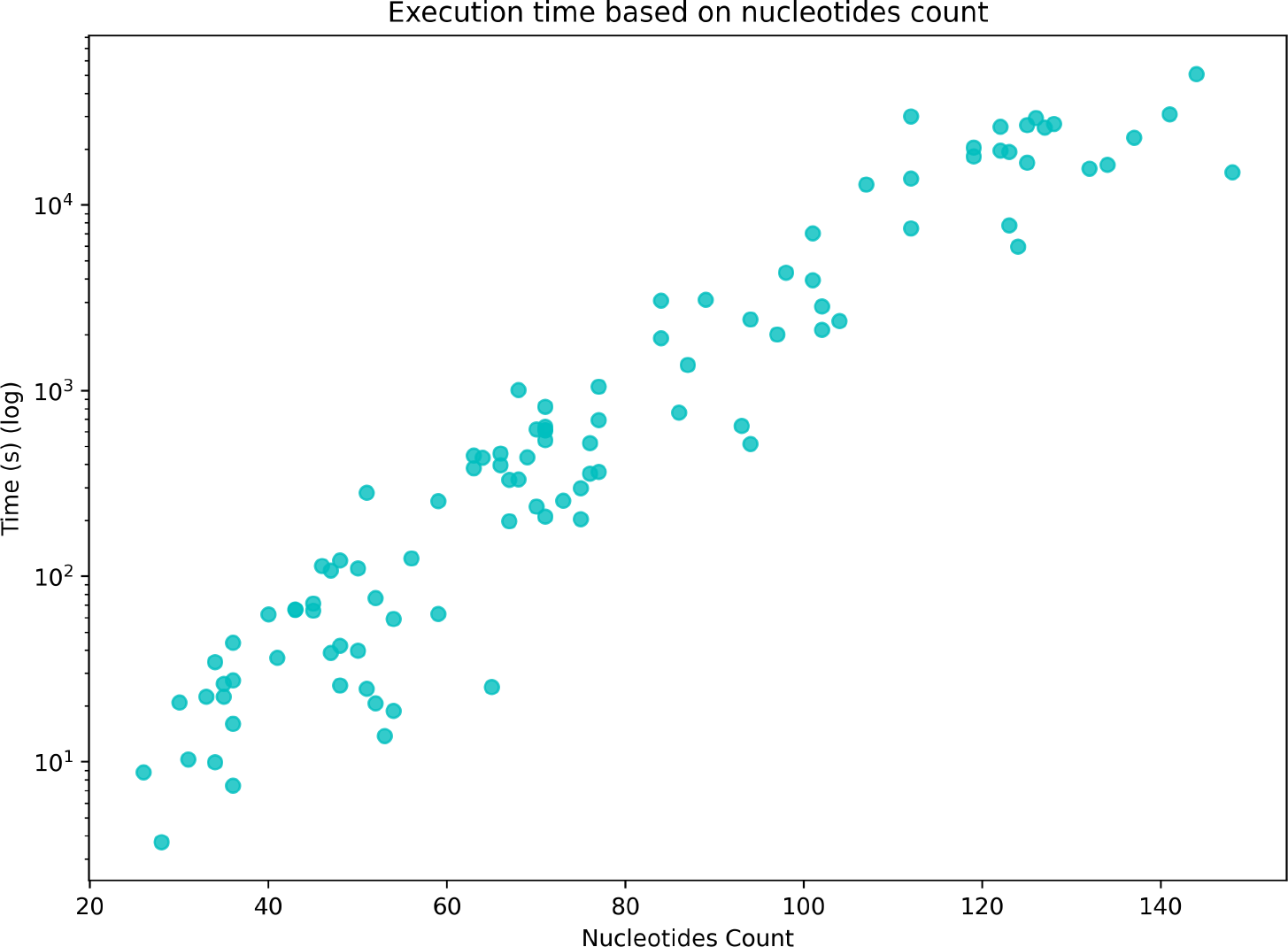
Execution Time-based on nucleotide count of the sequence at. *α* = 0.1. A maximum of 10^4^s was allowed, and 14 sequences didn’t return a solution in that time.

